# Wild emmer contribution in wheat domestication and adaptation to new environments

**DOI:** 10.1101/2023.10.25.563983

**Authors:** Alice Iob, Laura Botigué

## Abstract

Wheat is a staple crop, and its production is heavily threatened by climate change and soil erosion. Breeding programmes commonly rely on the wild progenitor, wild emmer wheat, *Triticum turgidum* subsp. *dicoccoides*, to find markers associated with traits conferring higher resistance to biotic and abiotic stresses, albeit how its genetic diversity contributed into domestication and adaptation to agricultural systems has never been studied. We explore the population structure and the influence of wild emmer populations from both the Northern and Southern Levant during the domestication process. Additionally, we examine their potential contribution to facilitating the adaptation and dispersal of domestic landraces to new environments. We quantify the genomic proportion of wild Southern Levant ancestry in two different domestic germplasms, including landraces from Europe, Africa and Asia. We obtain direct evidence that as much as 26% of the genome has Southern Levant ancestry in the population from Europe, and up to 40% in the population from Africa and Asia. We also estimate the time since admixture of the two wild populations to produce the domestic forms, obtaining two dates, one matching the domestication in Southwest Asia (ca 9500 BP) and the other matching the dispersal towards Africa (ca 6500 BP). We also inquire about the possible adaptive role of wild emmer from the Southern Levant into domestication and dispersal and find an overrepresentation of genes associated with resistance to biotic stress and drought. Overall, our work provides more information on the origins of domestic wheat and highlights the potential of modern domestic landraces of emmer wheat in the study of the genetic basis of resilience. Modeling wheat genome evolution under different demographic scenarios is needed to confirm the observed signals of positive selection and facilitate the use of emmer landraces in future wheat breeding programmes.

## Introduction

Over the past decades human food supply has become increasingly dependent on the cultivation of a few elite cultivars from a handful of species, prioritizing yield and caloric content while often overlooking other essential traits. This breeding process has led to a decrease in genetic diversity (Khoury et al., 2022), including variation conferring resistance to biotic and abiotic stresses, making modern cultivars heavily dependent on intensive inputs of water, fertilizers and pesticides. Consequently, agriculture is one of the major contributors to ecological degradation at the global scale, encompassing biodiversity loss (Gonthier et al., 2014), soil erosion and climate change. At the same time, climate change threatens the resilience and stability of our food system (Streit Krug et al., 2023) and great concerns arise from the decrease in productivity and increase in vulnerability of agriculture. In the past years global agricultural productivity has decreased (Arora, 2019; Ortiz-Bobea et al., 2021), in part due to the lack of sufficient diversity in elite cultivars to adapt and maintain yields amidst climate change (Hufford et al., 2019; Labeyrie et al., 2021; Streit Krug et al., 2023). Furthermore, predictions of greenhouse gas (GHG) emissions under the current food system would prevent the achievement of the targeted 1.5°C or 2°C degrees limits, even if all non-food emissions were to rapidly decrease (Clark et al., 2020).

The scientific community has turned to crop wild relatives (CWR) and traditional landraces to face these challenges. They constitute reservoirs of genetic diversity, harboring variation that allows them to adapt to diverse climatic conditions, and to grow with little input. (Cortés & López-Hernández, 2021; Marone et al., 2021). To incorporate desirable variants into high-performing cultivars, it is essential to conduct phenotyping of CWR and traditional landraces, while simultaneously preserving and characterizing their genetic diversity. At the same time, understanding genome evolution during domestication and dispersal to new environments is essential to identify genomic regions under positive selection, find polymorphisms influencing traits of agronomic interest and ultimately contribute to the development of new breeding strategies after functional validation (Venkateswaran et al., 2019).

Wheat is one of the most widely cultivated crops on earth (FAO, 2021), and it is seriously threatened by climate change. Each degree-Celsius increase in global mean temperature is predicted to reduce global yields by 6.0% (C. Zhao et al., 2017), in part due to the extremely low genetic variability of modern cultivars. Recent investigations have revealed that only a small fraction of the extensive diversity present in its ancestor wild emmer (*Triticum turgidum* subsp. *dicoccoides*) is retained within its domestic tetraploid descendants, and only 1.1% in bread wheat cultivars (Z. Wang et al., 2022). Domesticated emmer wheat, the ancestor of bread wheat, is known to be one of the most ancient cultivated cereals and is characterized by qualities such as high fiber content, resistance to biotic and abiotic stresses and ability to grow in adverse environmental conditions (Lucas et al., 2017; Saleh, 2020). Despite the exciting potential of emmer wheat to contribute to wheat improvement (Mohammadi et al., 2021; Zaharieva et al., 2010; Zohary, 2013), the evolution from wild to domestic emmer forms and the differentiation of domestic emmer are still largely unexplored topics.

Wild emmer wheat only grows in Southwest Asia (the Fertile Crescent) and is formed by two distinct populations living in different environments: one in the Northern Levant, mountainous, cold and humid, and the other in the Southern Levant, close to the Mediterranean coast with milder and dryer weather (Ozkan et al., 2011). The origins of domestic wheat remain uncertain. Several studies have shown that domestic emmer is genetically closer to wild emmer form the Northern Levant, as evidenced by phylogenetic and genome-wide analyses (Avni et al., 2017; Luo et al., 2007; Ozkan et al., 2002), but multiple studies have demonstrated that wild emmer from Southern Levant also contributed to the domestic gene pool (Cheng et al., 2019; Iob & Botigué, 2022; Pont et al., 2019). The model currently accepted distinguishes between a first phase of intensive exploitation and increasing management of wild or proto-domestic emmer starting in the Southern Levant, followed by dispersal and hybridization in Northern Levant leading to the emergence of the fully domestic phenotype, centuries later (Arranz-Otaegui et al., 2016; Civáň et al., 2013; Oliveira et al., 2020; Z. Wang et al., 2022). Fully domestic emmer would then spread out of the Fertile Crescent, adapting to new environments (Avni et al., 2017; Maccaferri et al., 2019; Zaharieva et al., 2010). This model aligns with evidence supporting the contribution of the wild emmer from the Southern Levant population at least one haplotype associated with the non-brittle rachis (Nave et al., 2019). The loss of rachis brittleness is considered the quintessential trait in cereal domestication, as it disrupts the plant’s natural seed dispersal mechanism. It is determined by two recessive mutations, one in chromosome 3A (*Tt-BRT1-A*) and the other in chromosome 3B (*Tt-BRT1-B*). According to this study, the domestic allele in chromosome 3A could be derived from the Northern Levant population, and the one in chromosome 3B from the Southern Levant one. However, a third scenario in which both alleles originated in Southern Levant could not be ruled out.

Despite these advancements, several key questions are still unanswered, including the timing and composition of wild admixture, as well as the extent of the Southern Levant’s contribution to the domesticated emmer gene pool. Additionally, some researchers continue to consider the genetic similarities between domestic emmer and its wild relatives from the Northern Levant as support for the formerly popular hypothesis that domestic emmer descended monophyletically from this wild population (X. Zhao et al., 2023).

Less attention has been given to the process of adaptation during the expansion of crops from their centers of origin (Janzen et al., 2019). Domestic emmer wheat spread from the Fertile Crescent to the Balkans, Western Europe, the Mediterranean basin, Eastern Africa, and eventually reached India adapting along the way to a wide range of different ecosystems (Maccaferri et al., 2019; Zaharieva et al., 2010). Adaptation can occur *de novo,* on standing variation or, when introduced through gene flow from a wild population, through a process known as *adaptive introgression*. Adaptive introgression has been proposed to have enabled changes in flowering time in flax during its expansion into Central Europe (Gutaker et al., 2019) or in fish living at high altitude (Qian et al., 2023).

Previous studies on the population structure of emmer wheat (Iob & Botigué, 2022), including wild and domestic specimens, identified two differentiated domestic germplasms, one representing the northwestern route of dispersal (DNW) into the Caucasus and Europe and another reflecting the southeastern route of dispersal (DSE) into Africa and Asia. Furthermore, evidence of extensive gene flow from wild emmer from Southern Levant into the DSE population was detected, explaining the differentiation between the two domestic germplasms. The uneven contribution from the wild Southern Levant population can be explained by two plausible scenarios: genetic drift following a reticulated domestication event, or hybridization during domestic wheat dispersal into Africa.

In this study, we aim to quantify the contribution of the wild emmer population from Southern Levant in the domestic pool, model the time since admixture and determine whether gene flow was linked to positive selection. Using haplotype-based techniques and selection statistics, we aim to gain insights into adaptive traits in wild emmer wheat and landraces, which can lead to the identification of advantageous alleles in front of the challenges of climate change in wheat cultivation. The identification of genomic regions that received one or the other influx from wild Southern Levant population, can aid understanding domestication mechanisms as well as adaptation.

## Results

Earlier research on the population structure of wild and domestic emmer wheat revealed that domestic emmer germplasm is generally more closely related to wild emmer samples from the Northern Levant. However, there was evidence of gene flow from the wild emmer population in the Southern Levant to the domestic germplasm from Africa and India (Iob & Botigué, 2022; Scott et al., 2019). To further characterize the contribution of the genetically diverse wild emmer from the Southern Levant to the domestic populations, we analyzed the genetic variability of an emmer collection (Zhou et al., 2020) containing specimens from Europe and Africa (Figure 1A and S. Table 1).

**Figure 1:**
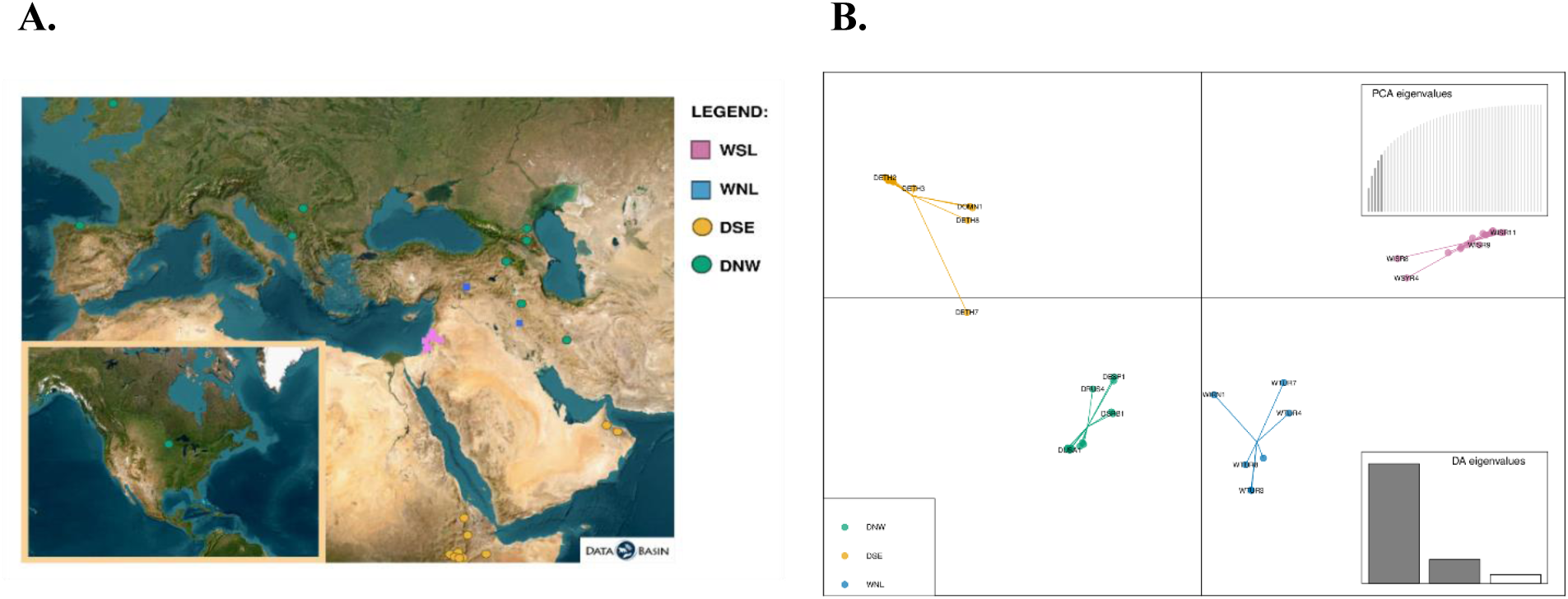
**1A**: Distribution of sites of the samples analysed in this study, colored according to the genetic clustering identified in the analyses. WSL: Wild Southern Levant, WNL: Wild Northern Levant, DNW: domestic northwest route of dispersal, DSE: domestic southeast route of dispersal. Map was created in DataBasin (https://databasin.org/). **1B**: Discriminant Analysis of Principal Components (DAPC) retaining 5 PC and 2 DA. The relationships between samples are the same as in Iob & Botigué 2022.

We applied a strict filtering approach to remove all potential artifacts from our dataset (see Methods). Specifically, after quality filtering we removed all heterozygote sites to eliminate artificial polymorphisms arising from the misalignment of repetitive regions, which reduced the number of SNPs by almost a half (from 66M to ca 38M, representing the “unmasked dataset”), and we further masked low complexity and repetitive regions based on the reference genome. This led to the retention of ca 10M SNPs representing the “masked dataset”. Given that this approach drastically reduced the number of SNPs for analysis, we tested the effect of the filtering by performing a Discriminant Analysis of Principal Components (DAPC), which confirms previously identified patterns. Wild samples from Southern Levant (Israel, Syria and Lebanon) cluster together and we henceforth refer to them as WSL population, while wild samples from Northern Levant (Turkey and Iran) form another cluster that we refer to as WNL. Domestic samples are distributed in two clusters, one containing specimens from Ethiopia and Oman, what we refer to as the DSE population, and another containing samples from Europe, the Caucasus, the Balkans and Iran, forming what we refer to as the DNW population. As previously assessed, the WNL population is closer to the domestic populations than WSL, and closest to the DNW group, while DSE looks more differentiated (Figure 1B). These results, that follow those obtained in Iob and Botigué 2022, show that the removal of heterozygotes and the masking of low complexity regions do not affect the relationships between populations within the dataset, while it removes potential unreliable signals.

### Wild ancestry of domestic populations

The use of unsupervised clustering algorithms to investigate the structure of the emmer wheat populations does not allow to model the contribution of each wild population in the domestic germplasm. The high degree of self-pollination in wheat combined with the low recombination rates in pericentromeric regions are translated in low levels of genome diversity. Such reduced diversity is interpreted as little admixture proportions at the genome level by these algorithms, as previously observed (Iob & Botigué, 2022). We used SourceFind, a haplotype-based method, to assess the extent of each wild population’s contribution to the two domestic populations, DNW and DSE.

Variability in both domestic populations arises mostly from the WNL group, in line with the literature (Avni et al., 2017; Oliveira et al., 2020). Nevertheless, we find that WSL not only contributed to the DSE domestic germplasm, but also to the genetic architecture of the DNW germplasm, obtaining direct genomic evidence of the influence of wild emmer from Southern Levant into the whole domestic pool. The estimated proportion of ancestry coming from WSL amounts to 26% for DNW and reaches as much as 40% of the whole genome contribution for DSE (Figures 2A and 2B).

**Fig. 2:**
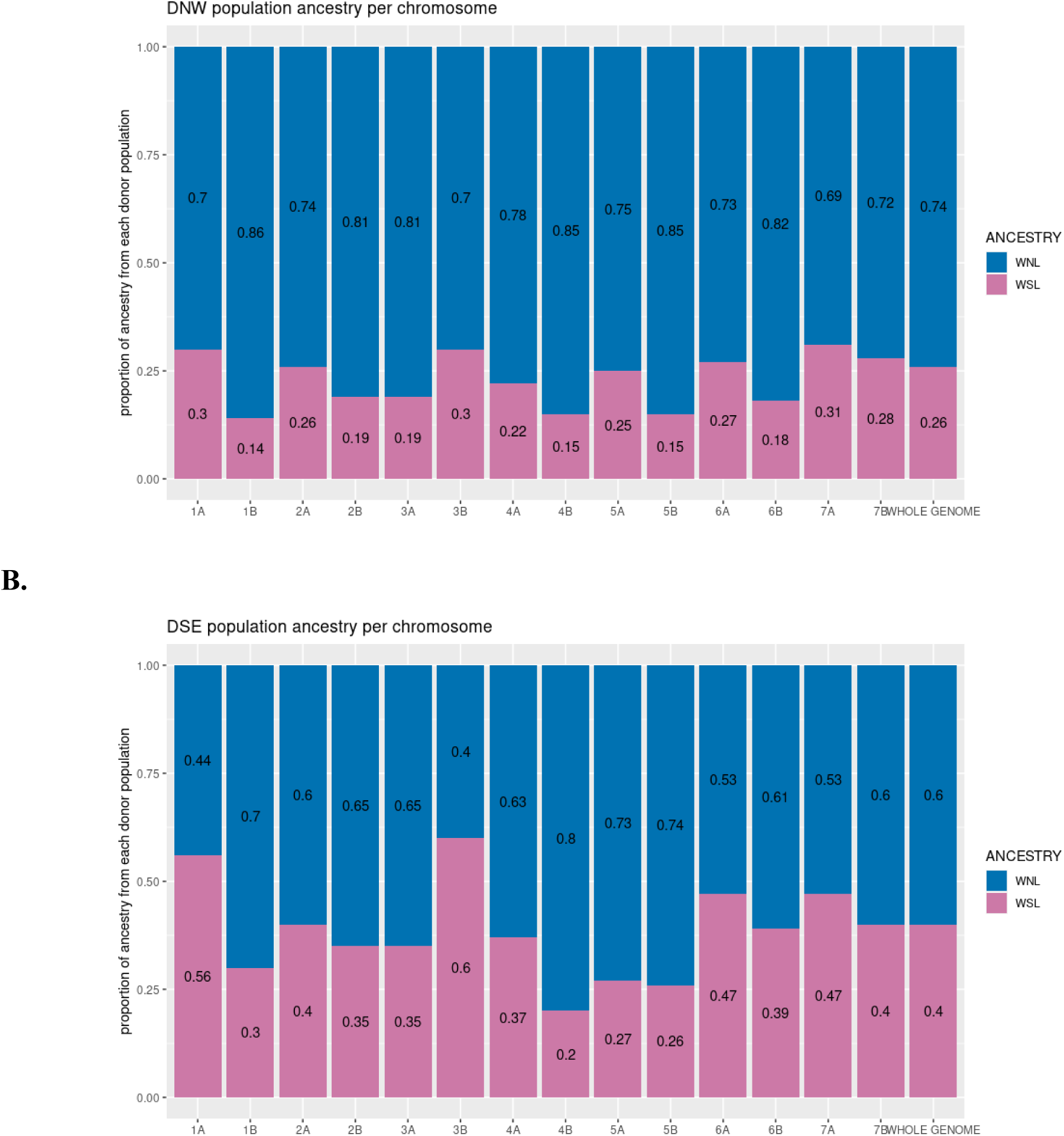
Wild ancestry in domestic populations as estimated by SourceFind. Results are reported for each chromosome and for the whole genome. **2A**: DNW; **2B**: DSE.

Results at the chromosome level show variable proportions of ancestry between chromosomes, which is expected by the combination of the effects of drift and differences in recombination rate along the chromosome. However, patterns of variation between chromosomes are shared by the two domestic populations. For instance, chromosomes 1B, 4B, 5A and 5B show the lowest levels of WSL ancestry in both populations, ranging from 14% to 26%, while chromosomes 1A and 3B, on the other hand, have the highest input from WSL in both populations, ranging from 30% to 60%. Interestingly, the copy of the *Tt-BTR1* gene responsible for the non-brittle rachis located in chromosome 3B has southern Levant ancestry (Nave et al., 2019), and out of the two gene copies is the one displaying the largest selective sweep (Scott et al., 2019), which is in line with the overall high proportion of Southern Levant ancestry in this chromosome. The presence of shared patterns of ancestry between the two domestic populations indicates that the contribution of WSL pre-dates the split between DSE and DNW.

We used the identified ancestry patterns to model the time since admixture between the two wild populations giving rise to the first domestic forms using FastGlobetrotter (Figure 3). When we considered DNW as the target population, we got one admixture event between WNL and WSL, dated 9,537 years ago (90% CI 4,900 – 13,900 years ago). This estimate is remarkably in line with the first archaeobotanical findings of fully domestic emmer (around 10,000 years ago) (Arranz-Otaegui et al., 2016). The majority contributing source is WNL, with 80% of ancestry, while WSL is the minority contributing source, with 20% ancestry. Interestingly, when we considered DSE as target population, the inferred date since admixture is 6,485 years ago (90% CI: 1,700 – 10,505 years ago), with WNL contributing 39% of the ancestry and WSL contributing 61%. This admixture date is compatible with the early dispersal of domestic emmer into the south, reaching Egypt between 7,500 and 6,500 years ago (Scott et al., 2019; Zaharieva et al., 2010).

**Fig 3:**
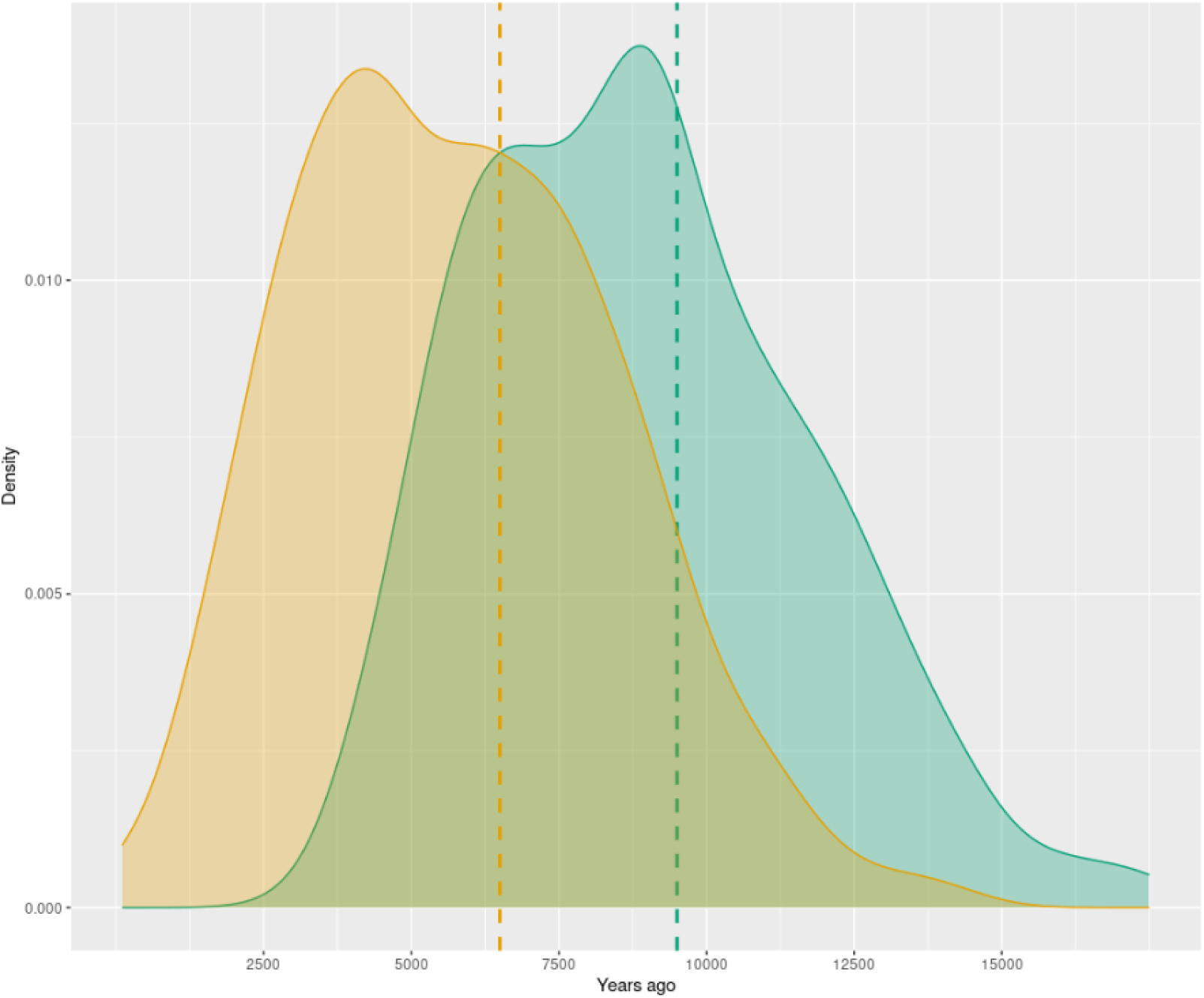
Density plot for the admixture dates estimates after 500 bootstrap iterations of Globetrotter. The x-axis shows the date since admixture (years ago). The green curve represents DNW, the yellow curve represents DSE.

These results support the hypothesis that the excess of WSL ancestry in modern DSE landraces comes from a hybridization event during the dispersal of early domestic forms into Africa.

Once we studied the wild ancestry proportions at the chromosome level, we aimed to study the contribution of wild population into the domestic pool at a finer scale. To do so, we calculated the absolute nucleotide divergence (Dxy) between populations along genomic intervals of 2Mb. As Dxy is not dependent on within-population diversity, it is better suited for the study of inbred populations. Averaging the values to get an estimate at the whole genome level, we obtained: Dxy WSL-DSE = 0.231, SD=0.066; Dxy WSL-DNW = 0.245, SD=0.066; Dxy WNL-DSE = 0.170, SD=0.093; Dxy WNL-DNW=0.150, SD= 0.081, confirming known relationships between the populations. Next, we compared the closeness of each domestic population to the two wild populations by plotting the joint distribution of Dxy scores between each domestic and the two wild populations (Figure 4A and 4B). In line with the haplotype-based results, most of the windows show that the two domestic populations have lower Dxy values (hence lower differentiation) with WNL than with WSL. However, differences can be observed between the two domestic populations. The DSE has an excess of WSL ancestry compared to DNW (1912 and 1015 out of 9881 windows showing lower Dxy values between DSE-WSL and DNW-WSL, respectively), Figure 4. This represents almost a two-fold increase, the same proportion that SourceFind estimated. On the contrary, DNW has more windows that show a high differentiation with WSL and low differentiation with WNL.

**Fig 4:**
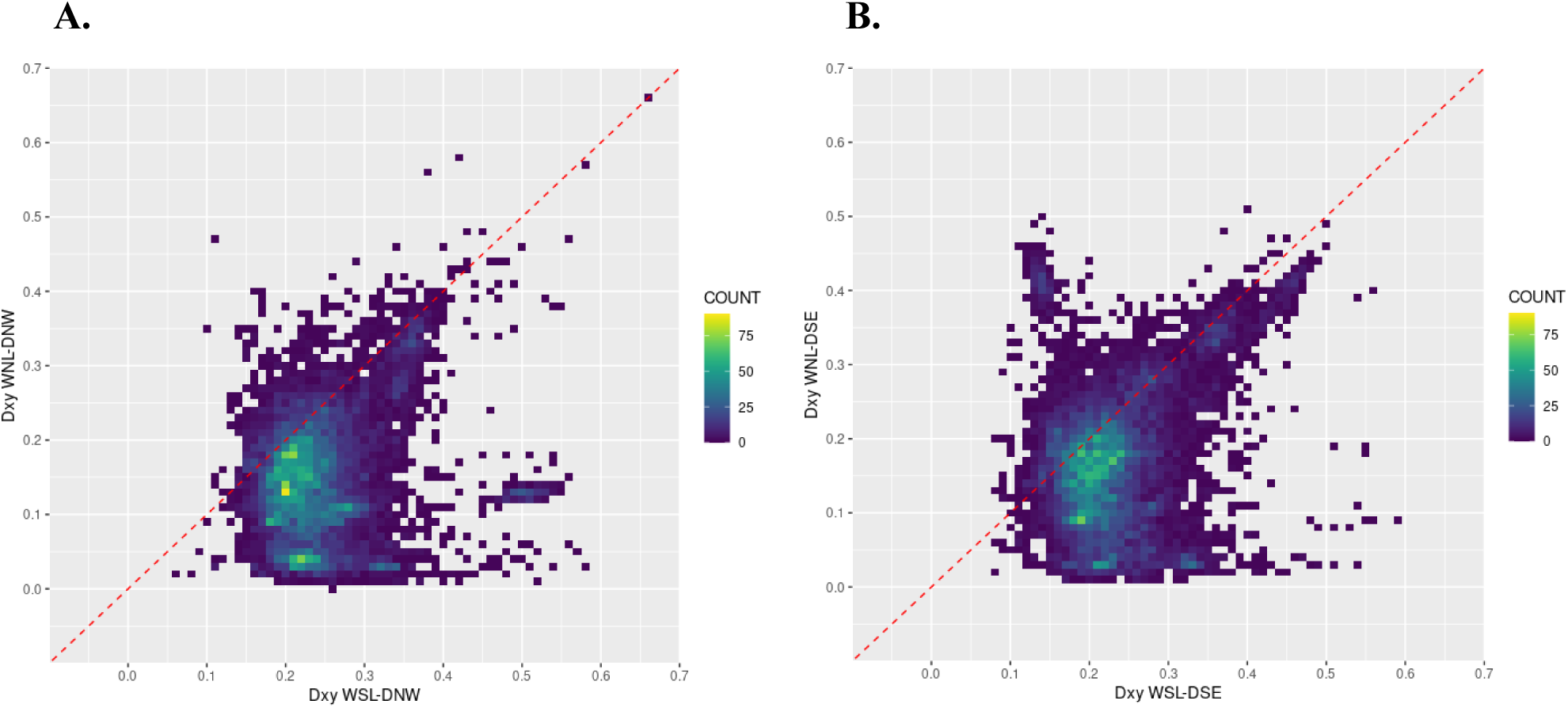
Dxy heatmap. X axis shows values of Dxy WSL-domestic and the y axis shows values of Dxy WNL-domestic. Each square represents a couple of values of Dxy for 2Mb genomic windows, indicating the relationship of such windows to one and the other wild population. The color depends on the number of occurrences of such Dxy value couples. The diagonal red line indicates equidistance to the two wild populations; values below such line indicate closer relatedness to the WNL population, while values above the diagonal line indicate closer relatedness to the WSL population. **4A**: DNW; **4B**: DSE, showing an excess of windows above the diagonal.

### Identification of shared WSL contribution to the domestic gene pool

In light of the results obtained, we propose a two-phase contribution of WSL to the domestic pool. An initial phase in which WSL hybridized with WNL (probably in the Northern Levant region in light of archaeobotanical evidence) to generate the domestic wheat, and a second phase in which WSL hybridized with the already domestic wheat spreading into Africa.

The legacy of wild populations from the Southern Levant in modern emmer wheat landraces has only been studied with regards to the evolution of the brittle rachis trait, but it has never been investigated at the whole genome level. We first focused on the influence of WSL in the domestication process by identifying regions of the genome where the two domestics show low differentiation with WSL and are genetically similar between them. For all those windows showing lower Dxy values between domestics and WSL than between domestic and WNL, we selected those that had a Dxy score between the two domestics smaller than 0.05. This pattern of differentiation reflects the contribution of the WSL into the domestic pool that has remained relatively unchanged in the two domestic germplasms since the domestication process.

This led to the identification of 184 2-Mb, overlapping windows (Table S2), corresponding to a total of 279Mb of Southern Levant origin that remain similar in the two domestic populations. Notably, the windows containing the *Tt-BTR1-B* locus controlling rachis brittleness on chromosome 3B (of putative Southern Levant origin in the domestic forms) were included (96-98 Mb in chromosome 3B). A similar pattern was observed when genetic variability in the domestics was compared with that of WSL. XP-EHH score was in the top 5 and top 1 percentile for DSE and DNW, respectively in the 94-96Mb window, but not in the 96-98Mb. These results reflect the importance of using overlapping windows and support the idea that using a genetic distance statistic such as Dxy it is possible to retrieve the signal of positive selection. On the other hand, the window containing the *Tt-BRT1-A* locus on chromosome 3A shows lower distance to the WNL population for both domestics (Dxy WSL-domestics 0,231; Dxy WNL-domestics 0,173, Dxy DSE-DNW 0,007). These results support one of the two-step domestication scenarios hypothesized by Nave et al. 2019, namely an independent origin of the domestic *Tt-BTR1* genes in north and south Levant and subsequent hybridization that would generate the fully domestic phenotype.

Within the 184 overlapping windows, we found 170 SNPs with a predicted high impact on the biological function of the protein products in 85 genes, and over 4500 (4535) with moderate impact on the biological function in 829 genes (Table S3). Three genes carrying high impact variants drive the statistical overrepresentation in *systemic acquired resistance*, (GO:0009627, adjusted P-value 0,0212), a mechanism of induced defense occurring in the distal parts of the plant following localized infection and conferring protection against a broad spectrum of microorganisms (Durrant & Dong, 2004). Interestingly, the canonical Ensembl transcript of one of these genes has an NPR3 domain, where the high impact polymorphism is located. It is possible that two transcripts from different genes have been fused, one of them coding for an NPR3 protein. Only NPR1 proteins have been annotated in this reference genome, so it is likely that this fused transcript contains an NPR3 protein. The other two genes encode for Lipid Transfer Proteins that have also been associated with plant immunity (Finkina et al., 2016).

The genes affected by moderate and high impact variants taken together show enrichment in galactoside 2-alpha-L-fucosyltransferase activity (GO:0008107, adjusted P-value 0,0237), alpha-(1,2)-fucosyltransferase activity (GO:0031127, adjusted P-value 0,00237), and binding (GO:0005488, adjusted P-value 0,0010), molecular functions known to be related to the synthesis of cell wall matrix and hence involved in several biological processes (Reiter, 2002; Tryfona et al., 2014). Fucosyl transferases are known to play a role in cell wall biosynthesis in cereals (Hazen et al., 2003) and their activity has been linked to both salt sensitivity (Tryfona et al., 2014) and immunity (Zhang et al., 2019).

### Introgression from WSL to DSE

We next focused on the unique contribution of WSL to the DSE population, which we hypothesize is the consequence of the second hybridization phase during the early dispersal of the first domestic wheat forms out of the Fertile Crescent. We identified those regions of the emmer wheat genome in which not only DSE is closer to WSL than to WNL but also the differentiation between the two domestic populations is high. From the calculation of Dxy as described above, we selected only those windows in which Dxy WSL-DSE is smaller than Dxy WNL-DSE and Dxy DSE-DNW is bigger than 0.33, based on patterns of Dxy (top 5% values).

The analysis revealed 247 2-Mb overlapping windows, corresponding to 271Mb in total, that show contribution from WSL to DSE only (Table S4). Within the 247 windows, we found 311 SNPs that have high (16 SNPs) or moderate impact on protein products of 186 genes (Table S5). Such genes showed overrepresentation for molecular functions, biological processes and cellular components mainly related to cell cycle, metabolism, cellular organization, membrane and organelles, as reported in table 1.

**Table 1:** overrepresentation test results for genes related to introgression from WSL to DSE.

| SOURCE | TERM NAME | TERM ID | ADJUSTED P-VALUE |
| --- | --- | --- | --- |
| GO:MF | molecular_function | GO:0003674 | 0,0021 |
| GO:BP | cell cycle G1/S phase transition | GO:0044843 | 0,0011 |
| GO:BP | G1/S transition of mitotic cell cycle | GO:0000082 | 0,0011 |
| GO:BP | metabolic process | GO:0008152 | 0,0045 |
| GO:BP | biological_process | GO:0008150 | 0,0082 |
| GO:BP | primary metabolic process | GO:0044238 | 0,0162 |
| GO:BP | organic substance metabolic process | GO:0071704 | 0,0172 |
| GO:BP | intracellular transport | GO:0046907 | 0,0455 |
| GO:BP | establishment of localization in cell | GO:0051649 | 0,0488 |
| GO:CC | membrane coat | GO:0030117 | 0,0001 |
| GO:CC | coated membrane | GO:0048475 | 0,0001 |
| GO:CC | cytoplasm | GO:0005737 | 0,0010 |
| GO:CC | intracellular anatomical structure | GO:0005622 | 0,0028 |
| GO:CC | membrane protein complex | GO:0098796 | 0,0067 |
| GO:CC | cellular_component | GO:0005575 | 0,0080 |
| GO:CC | cellular anatomical entity | GO:0110165 | 0,0173 |
| GO:CC | chloroplast | GO:0009507 | 0,0319 |
| GO:CC | intracellular membrane-bounded organelle | GO:0043231 | 0,0334 |
| GO:CC | membrane-bounded organelle | GO:0043227 | 0,0339 |
| GO:CC | intracellular organelle | GO:0043229 | 0,0354 |
| GO:CC | organelle | GO:0043226 | 0,0358 |
| GO:CC | plastid | GO:0009536 | 0,0396 |

Forty percent of these windows (n=99) are found in a contiguous stretch in chromosome 4B, between 189 and 378 Mb (Figure 5A). The low recombination rate in this area, 8.6 * 10^−9^ cM/bp, based on Maccaferri 2019, explains the length of the region. Within this region the two domestic populations are quite divergent (Dxy=0.4356, SD=0.0236), and DNW is clearly closer to WNL, as shown by Dxy values (Dxy WNL-DNW=0.1288, Dxy WSL-DNW=0.5027) (Fig. 5B), evidencing that this region has different wild ancestries in the two domestic populations. These results are surprising, since chromosome 4B is among the chromosomes with fewest WSL ancestry in both domestic populations (15% in DNW and 20% in DSE) according to SourceFind. This slight difference of 5% in the estimated contribution of WSL between the two domestics rules out that the signal in chromosome 4B is related with the second hybridization phase. By examining Dxy values for DNW and DSE with the two wild populations we could corroborate that for most of the chromosome not only were both domestics closer to WNL than WSL, but that Dxy values were low, ranging between 0 and 0.1. Rather than adaptive introgression, these results support an evolutionary constraint to preserve ancestry from WNL during the domestication process and that only the centromeric region in DSE accumulated WSL ancestry. This hypothesis is further supported by the fact that according to SourceFind, only chromosome 4B and chromosome 5A have similar amounts of WSL ancestry in the two domestics, while other chromosomes display between 11 and 20% increase in WSL ancestry in DSE compared to DNW. Notably, chromosome 5A harbors the Q gene, also called the domestication gene for its many effects on different phenotypes associated with early crop improvement (Simons et al., 2006).

**Figure 5:**
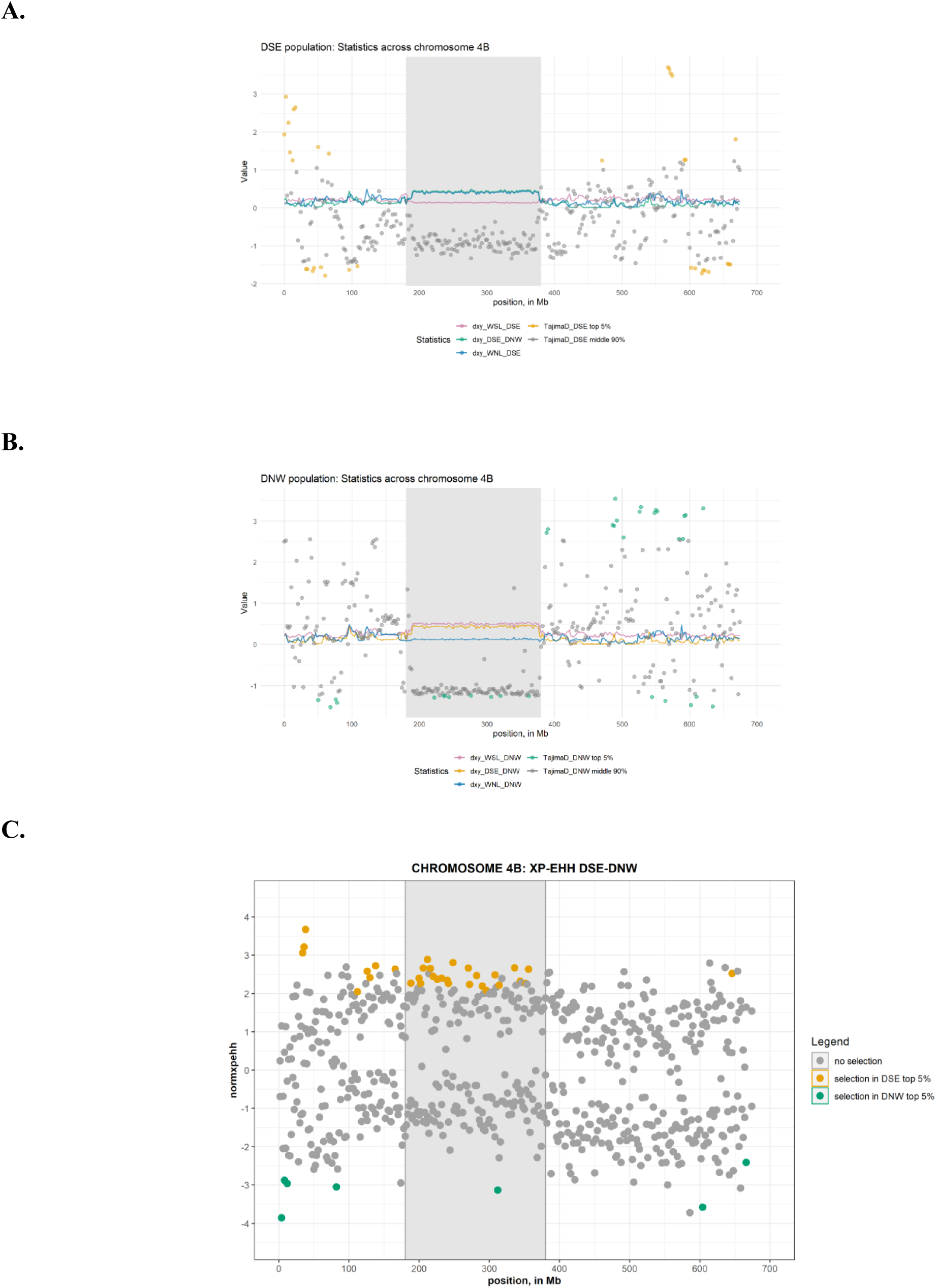
statistics across chromosome 4B. Tajima’s D and XPEHH are calculated in 2Mb non-overlapping windows, while Dxy in 2Mb overlapping windows, with a step of 1Mb. **5A**: Taijma’s D and Dxy for DSE population. Yellow points show top 5% Tajima’s D values; **5B**: Taijma’s D and Dxy for DNW population. Green points show the top 5% Tajima’s D values; **5C**: XP-EHH DSE-DNW. Yellow points show top 5% values for selection in DSE, green points show top 5% values for selection in DNW

The WSL ancestry stretch in DSE could have evolved through the effect of drift or selection. Since our goal was to assess the role of wild wheat from the Southern Levant in the domestic pool, we next investigated this region for signals of selection. We calculated Tajima’s D scores and found that both domestic populations have negative Tajima’s D values, especially DNW (Figs 5A and 5B). On the other hand, DSE shows lower nucleotide diversity (PI=0.0333, SD=0.0036) than DNW (PI=0.1012, SD=0.0109). Both negative Tajima’s D and reduced nucleotide diversity are potential indicators of a selective sweep. However, this excess of rare alleles (Tajima’s D < 0) can also be explained by the combination of inbreeding and low recombination rate of pericentromeric regions in wheat. For this reason, we decided to compute the XP-EHH statistic in DNW vs. DSE for the identification of a selection signal within this region. XP-EHH is a haplotype-based statistic that allows for the identification of recent selective sweeps. XP-EHH implicitly accounts for the recombination rate within the genome and is also less sensitive to population structure, as it detects selection by comparing haplotypes between two populations (Eydivandi et al., 2021). XP-EHH results (Fig 5C) show top values for DSE within this region, indicating strong evidence of a selective sweep in this population.

We found 71 genes affected by moderate or high impact variants in this region. These genes show enrichment in organic substance metabolic process (GO:077170, adjusted p-value 0,038), chloroplast (GO:0009507, adjusted p-value 0,001), plastid (GO:0009536, adjusted p-value 0,001), intracellular membrane bounded organelle (GO:0043231, adjusted p-value 0,032) and membrane-bounded organelle (GO:0043227, adjusted p-value 0,033). Plant metabolism plays a crucial role in acclimation and survival under stress conditions (Fraire-Velazquez & Emmanuel, 2013), and the chloroplast is considered a metabolic center with key role in adaptation to heat stress (Q. L. Wang et al., 2018).

Overall, these results suggest that the WSL population may have played a role in adaptation of domestic wheat to the African milder and dryer climates.

In addition to the region in chromosome 4B, the differentiation-based filtering also led to the identification of a smaller stretch of 2-Mb windows in chromosome 6A, between 177 and 208 Mb. Windows within this region also produce negative Tajima’s D scores. Within this region, the XP-EHH values don’t reach the top 5% for DSE, but the values are higher than the chromosome average (average within this region =1.48; chromosome average = 1.2).

These results are compatible with a selective process in wheat from DSE in genomic regions with a WSL genetic ancestry. Moreover, these results highlight once again the necessity of applying a combination of different methods when analyzing introgression and selection.

## Discussion

Emmer wheat, nowadays considered as a “neglected” crop, is cultivated only in a few areas of the world such as India, Yemen or Ethiopia, where it is consumed in moderate amounts (Zaharieva et al., 2010). Its wild progenitor, wild emmer wheat, has been used as a genetic resource for QTL mapping and potential wheat improvement. However, the generation of wild and domestic wheat hybrids carry many unfavorable traits, making the task towards wheat improvement slow and arduous (Engels & Thormann, 2020). On the contrary, modern landraces of domestic emmer carry many traits of interest for wheat improvement without the burden of undesirable wild traits (Swarup et al., 2021). At the genome level, domestic emmer has been often investigated in the context of genetic variation in durum and bread wheat, while its individual genetic characteristics have received substantially less attention.

Our analysis based on genetic differentiation and polymorphisms with a probable high impact on the biological function reveal that several regions of the genome with a most likely ancestry from Southern Levant may have been selected both during domestication and during the early dispersal of the domestic forms into Africa. Interestingly, our results highlight the potential of emmer wheat landraces in wheat improvement. When focusing on selection during the domestication process, we found a significant overrepresentation in genes involved in systemic acquired resistance (SAR, GO:0009627) in the regions of the genome with shared WSL ancestry in the two domestic populations. Such enrichment holds great scientific interest, as plant resistance to herbivory and pathogens is a primary phenotype found predominantly in wild ancestral species (Carmona et al., 2011; Chaudhary, 2013; Y. H. Chen et al., 2015), and typically depleted in modern wheat cultivars. Wild plants, constantly exposed to diverse pathogens, rely on inherent genetic resistance for fitness and survival in natural habitats. However, in cultivated environments, the use of agronomic practices and chemical interventions gradually diminished the need for natural pathogen immunity in cultivated plants (Singh & van der Knaap, 2022). These results support the hypothesis that alleles increasing biological resistance coming from wild wheat from Southern Levant would have been selected and conserved in traditional emmer landraces but lost in elite cultivars of modern bread and durum wheat, as reflected by efforts to introduce resistance genes from their CWR (Hajjar & Hodgkin, 2007).

Haplotype-based analysis carried out with sourceFind allowed us not only to quantify the amount of wild Southern Levant ancestry in domestic populations but also detect the post-domestication hybridization event in domestic emmer landraces from Africa and the Arabian Peninsula, previously detected at a lower resolution. Remarkably, genes containing polymorphisms with an estimated high and moderate impact in the biological function were overrepresented in processes that have been associated to drought stress and tolerance in maize by more than one study (Jiang et al., 2014; Xu et al., 2014). Among the overrepresented cellular components, it is worth highlighting organelles and plastids. A recent study on drought resistance in wheat (Lv et al., 2020) suggest that that cellular organizations play a crucial role in drought stress, highlighting the susceptibility of the cytoplasm, peroxisome, and chloroplast to drought stress treatment during early developmental stages in wheat. The authors also point out that drought stress significantly affects various processes, including, among others, the cell cycle. Chloroplast overrepresentation largely stems from the long stretch in chromosome 4B. Incidentally, recent studies have shown that the chloroplast is strongly affected by heat stress (Akter & Rafiqul Islam, 2017) and it is known to undergo metabolic reprogramming in response to it (Hu et al., 2020; Q. L. Wang et al., 2018). Moreover, one of the genes in this region that is involved in all categories showing enrichment, is *TRITD4BV1G106550*, which codes for Photosystem II Psb27 protein, a protein that transiently binds to PSII assembly intermediates before a fully functional PSII is formed (Liu et al., 2011). It has been shown that such protein is involved in response to photodamage of PSII in *Arabidopsis* (H. Chen et al., 2006) and a recent study showed that Psb27 participates in light energy dissipation to allow correct maturation of PSII (Johnson et al., 2022).

Overall, our results strongly suggest that there has been positive selection on regions of the genome of WSL ancestry in domestic wheat spreading towards Africa. This selection was likely linked to light, heat and drought stresses, and therefore adaptation to hotter and dryer climates. Modern emmer from these regions should be further investigated to validate the effect of these polymorphisms on heat and drought resistance.

More studies are needed to confirm that these sites are under positive selection indeed, both at the functional level and by comparing wheat genome evolution under drift and under adaptation. The combination of long chromosomes and self-pollination increases the effect of drift, but at the same time under certain conditions can facilitate the fixation of positive alleles. Looking for evidence of differential selective pressure in the two domestics, we did find overrepresentation of biological processes like transmembrane transport activity in DNW, and genes associated to 1,4-alpha-glucan branching enzyme activity (GO:0003844) in DSE, known to play a fundamental role in starch biosynthesis (Bahaji et al., 2014; Tetlow & Emes, 2014). However, genetic variability between the two domestics is substantially smaller than between the domestics and wild wheat from Southern Levant, the focus of this manuscript, and other methodological approaches are needed to confirm this potential evidence of selection.

### Refined insights into wheat domestication and dispersal

With regards to the historical significance, our results not only support a reticulated model of wheat domestication but also quantify the contribution of each wild population in the domestic pool, also at the chromosome level. We find that the proportion of WSL ancestry in the two domestics is correlated for most of the chromosomes, pointing to an initial phase of admixture that pre-dates the split of the domestic gene pool into the two populations. The proportions of each wild ancestry that we find suggest that the two proto-domestic populations hybridized in the Northern Levant, generating the fully domestic phenotype. This evidence is compatible with the presence of fully domestic archaeological assemblages in the NL starting from Middle-Late Pre-Pottery Neolithic B (MLPPB) and previous hypotheses (Oliveira et al., 2020). This genetic model also reconciles with archaeological models that indicate domestication as a long and patchy process, involving different wild populations and human communities (Asouti & Fuller, 2013; Fuller et al., 2023; Mithen et al., 2023).

Further evidence of the reticulated domestication is provided with our analyses on genetic differentiation and the *TtBTR* loci, which confirm a dual ancestry in chromosomes 3A and 3B. If we incorporate archaeobotanical evidence and previous results, the most plausible scenario is that the domestic allele would have emerged first in chromosome 3B in the Southern Levant, since it is the region where wild emmer was first managed during Pre-Pottery Neolithic A (PPNA) and the first findings of partially domestic wheat occur. During this period incipient agricultural communities in the Northern Levant were focused on other species, based on the archaeobotanical remains identified (Arranz-Otaegui et al., 2016). The second haplotype (on chromosome 3A) would have emerged later, when emmer cultivation increased in the Northern Levant. Admixture between the two proto-domestics carrying each of the two haplotypes would have led to the appearance of the fully domestic phenotype.

We also used haplotype information from the whole genome to estimate the time since admixture between the two wild populations to model a domestication scenario. The date obtained for DNW, 9500 years ago, is very close to the appearance of fully domestic assemblages in the archaeological record. For DSE, the time since the last major admixture event is around 6500 years ago, matching the Southern dispersal of emmer wheat to Africa (Özkan et al., 2011). Based on these results in combination with the other findings, we hypothesize that the additional 20% of WSL ancestry in DSE compared to DNW reflects a second wave of hybridization between the domestic forms and wild wheat from Southern Levant. Even if FastGlobetrotter indicates a single event of admixture, our results are limited by having only two source populations, which reduces the power of the model to distinguish between one or multiple hybridizations. Furthermore, emmer wheat has long chromosomes that rarely recombine around the pericentromeric regions forming long haplotypes.

In light of this, we propose a model of wheat dispersal towards Africa where hybridization occurred not between the wild populations in Northern and Southern Levant, but between the fully domestic forms spreading southwards from the Northern Levant and either wild populations from Southern Levant or, most likely, proto-domestic forms in the area. SourceFind and Dxy results support that the hybridization occurred predominantly on the Northern Levant genetic background, as reflected by the higher proportion of WNL-derived ancestry in both populations. A schematic representation of the main events leading to the appearance and diversification of the domestic landraces can be found in Figure 6.

**Figure 6:**
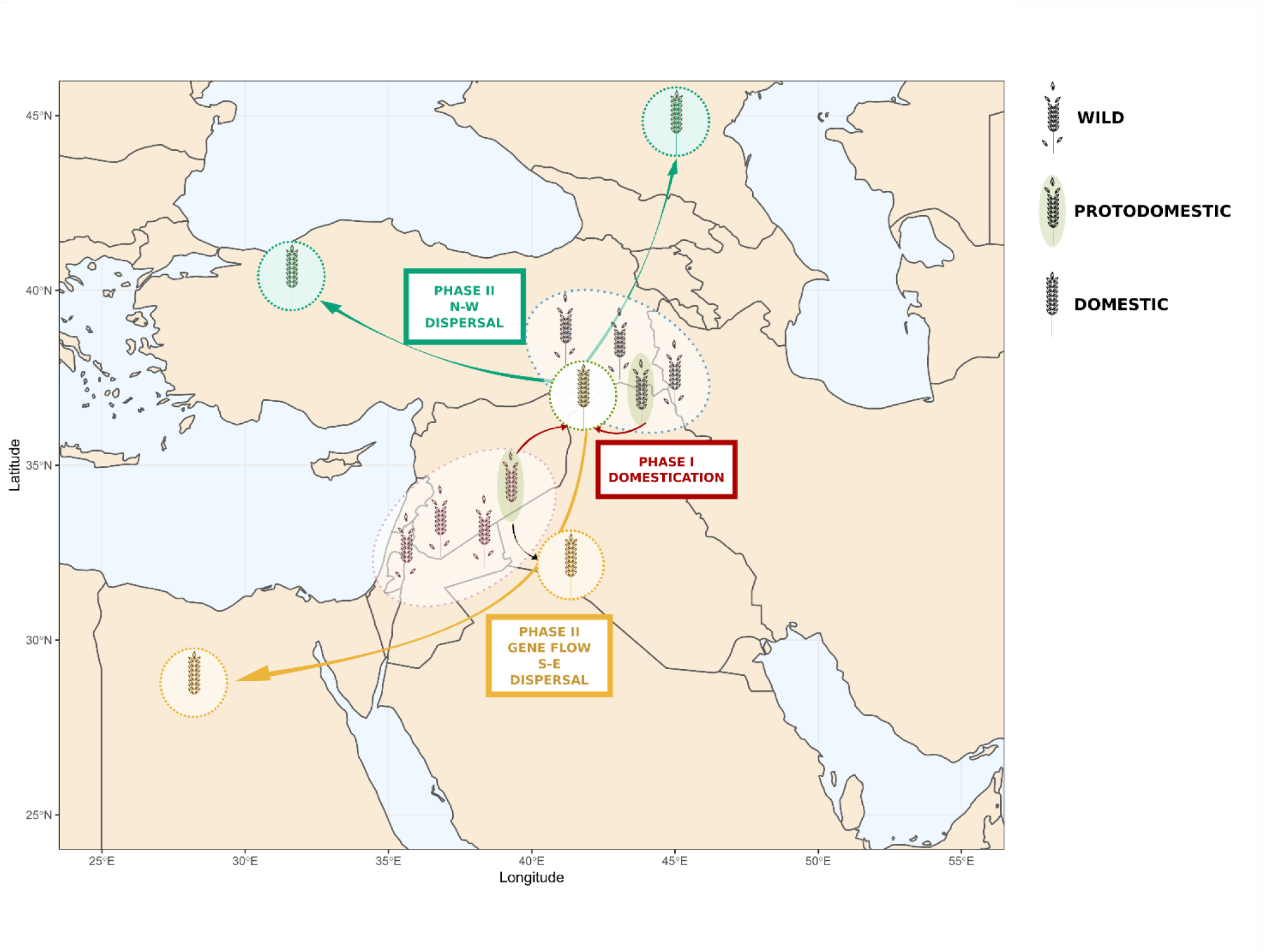
Schematic representation of the events leading to the appearance and diversification of domestic emmer. Wild, protodomestic and domestic status are represented through varying degrees of rachis brittleness. The pink and blue dotted circles represent the wild and protodomestic Southern and Northern Levant populations, respectively. The green and yellow circles represent the domestic NW and SE populations, respectively. In the first phase, protodomestic individuals from the SL and the NL hybridize (red arrows) in the north of SW-Asia (or Northern Levant), leading to the appearance of the domestic phenotype. In a second phase, domestic emmer that dispersed to the north and west of SW-Asia differentiated into contemporary DNW population (green arrows). Domestic emmer that dispersed to the south hybridized with wild or protodomestic emmer from the SL (black arrow), giving raise to the contemporary DSE population, that reached Africa and India (yellow arrow).

## Conclusions

Our genomic analysis of traditional emmer landraces and their wild relatives has unveiled new details of the domestication process. More importantly, our combined survey of ancestry and of genomic regions that show evidence of positive selection has revealed several aspects to take into consideration. First, studying different germplasms is crucial to have a comprehensive catalogue of genetic variations. This is true not only for traditional landraces, in this case with the inclusion of accessions from Ethiopia and Oman, but also Crop Wild Relatives. Generating a wheat pangenome would allow to better characterize genetic variability and include structural variation, giving another level of resolution of ancestral wheat genome evolution. Second, the lack of a good, parametrized demographic model for wheat does not allow us to distinguish between the effect of drift from that of adaptation. Despite this, our conservative approach has unveiled many regions that are candidates for positive selection and that have been previously associated with resistance to biotic and abiotic stresses. Efforts to understand wheat genome evolution under domestication and dispersal are essential to increase our ability to distinguish the genomic footprints of adaptation. Finally, our results show that not only CWR are a desirable tool for breeding improvement. Our understanding of how WSL contributed to the adaptation of domestic wheat suggests that these domestic landraces can serve as valuable resources for breeding, as they already possess a multitude of desirable traits.

## Materials and methods

In this study we analyze 55 emmer wheat samples, representative of two wild and two domestic populations: 7 are from Wild Northern Levant (WNL), 20 are from Wild Southern Levant (WSL), 15 are from the domestic southeastern route of dispersal (DSE) and 13 from the northwestern route of dispersal (DNW). Passport information is available in Supplementary Table 1 (Table S1). Samples were processed as in Iob and Botigué (2022), with further filtering applied. Briefly, reads were aligned to the durum reference genome (*Triticum turgidum* subsp. *durum,* Maccaferri et al., 2019) and variants were called in the whole dataset using GATK v. 4.1.6 (Van der Auwera, GA O’Connor, 2020). Hard filters were applied after genotype calling as in (Zhou et al., 2020), and 66M SNPs were kept. Emmer wheat is a highly homogeneous and repetitive genome, and as such is prone to read misalignment that translates into inflated (false) heterozygosity (Bukowski et al., 2018; Li & Wren, 2014). To circumvent this problem, we applied a very conservative approach and removed all sites that showed at least one heterozygote genotype. We refer to the resulting dataset as “unmasked dataset”, comprising ca 38 million (38 099 555) homozygote SNPs.

For all analyses involving the calculation of genetic differentiation and selection statistics, we masked SNPs located in low complexity and repetitive regions based on the masked version of the reference genome. To do so, we utilized the "generate_masked_ranges.py" script (Cook, D.E., GitHub), which translated the coordinates of masked regions into a bed file. Using this bed file, we excluded SNPs within these regions from the VCF file and ensured no missing information using the vcftools v.0.1.16 (Danecek et al., 2021) options "-exclude-bed" and "-max-missing 1." This filtered dataset, referred to as the "masked dataset", comprises over 10 million (10,398,941) SNPs. Additional filters specific to each analysis are discussed in their respective methodological sections.

To test the goodness of our strict filtering approach, we performed a DAPC (Discriminant Analysis of Principal Components) on the masked dataset. First, we filtered it for linkage disequilibrium, using plink v. 1.9 (Purcell et al., 2007), allowing a maximum -r^2 value of 0.1 calculated in 50 kb windows with a step size of 10 kb, reducing the dataset to ca 750 000 (748 812) variants. Then, we used the Adegenet package (R 4.1.0, R Core Team, 2021) to perform the DAPC, retaining 5 principal components (PC, based on the result of the xvalDapc function) and 2 discriminant analysis eigenvalues (DA).

### Wild Ancestry of domestic populations

In this study, we employed Chromopainter v.2 (Lawson et al., 2012), SourceFind v.2 (Chacón-Duque et al., 2018) and FastGlobetrotter (fastGT) (Wangkumhang et al., 2022), to estimate ancestry proportions and admixture times for a specific target population using source populations. We used this strategy to model the admixture process of the two domestic populations (DSE and DNW) separately, using the two wild populations (WSL and WNL) as sources. We followed the protocol described in (Hellenthal et al., 2014), and we utilized perl 5.26.0 and R 4.1.0 for its implementation. The unmasked dataset was phased and imputed using ShapeIT v.2 (Delaneau et al., 2014), applying the recombination rate of different chromosomic regions as published in Maccaferri et al., (2019). To avoid introduction of biases due to lack of heterozygote sites, we “haploydized” the dataset by removing one allele for each genotype, and ran all subsequent steps accordingly.

We first ran ChromoPainter to infer the proportion and number of haplotypes shared between individuals, using all samples as possible recipients but only wild samples as donors.

We performed an initial run to estimate switch rate parameters (-n) and global mutation parameters (-M) using ten iterations of expectation maximization per chromosome, and then we averaged the resulting values across all chromosomes, weighting by the number of SNPs per chromosome. The resulting fixed values were used to run ChromoPainter for all chromosomes and the final ancestry matrices (i.e., *.chuncklengths.out and *.chunckcounts.out files) were summed across chromosomes. We then ran ChromoPainter a second time, painting each domestic target using wild sources as donors only (disallowing "self-copying" from other members of the same population), to generate painting sample files (i.e. *samples.out files). The resulting copy vector files from the first ChromoPainter run, and the painting sample files from the second ChromoPainter run were used as input files for both SourceFind and FastGlobetrotter. We ran SourceFind using default parameters to infer the admixing proportion of WNL and WSL for both DSE and DNW.

In the fastGT runs (one for each domestic population), the null.ind parameter was set to 1, as recommended, to account for decay in linkage disequilibrium that may not be due to authentic admixture signals. We set the haploid option with “haploid.ind 1”, changed the curve range from 30 to 15 with "cr 15" to account for the slow decay of linkage in the emmer wheat genome, and the generation time to 1 year. We performed 500 bootstrap replicates and calculated the 90% confidence interval from the distributions of the replicates. FastGT assumes an outcrossing rate of 100%, but emmer wheat is a mostly self-pollinating species. For this reason, we adjusted the admixture times as follows: considering that emmer wheat has a selfing rate of around 99% (Dvořák, 2001; Golenberg, 1988), we expect an outcrossing event with a probability of 1%, or, in other words, 1 every 100 generations. Hence, we divided the estimate times since admixture by the outcrossing rate (Treal = Tinferred /(1/100)).

### Genetic differentiation

We calculated Dxy (absolute nucleotide divergence) to measure genetic distance between populations. We opted for Dxy over Fst because Dxy is unaffected by within-population diversity (Henderson & Brelsford, 2020). In self-pollinating species, diversity is generally lower than in outcrossing ones, and domestic populations tend to have reduced diversity compared to their wild relatives. These characteristics are crucial when comparing diversity levels and genetic distances between populations. While Fst relies on within-population genetic diversity and can yield high values when one population has low diversity in a specific genomic region, Dxy provides an absolute measure of genetic divergence between populations, making it a better choice for self-pollinating species in domestication studies. For the calculation of Dxy we used the masked dataset and a window size of 2 Mb and a step of 1Mb and applied the scripts from (Martin, S., GitHub): we converted vcf to “geno” format using parseVCF.py and then calculated Dxy using popgenWindows.py considering only windows with at least 100 good sites (options -w 2000000, -s 1000000, -m 100). We rounded the values of Dxy to two digits and we plotted the number of windows pairs with the same Dxy couples of values using ggplot2 (R 4.1.0).

### Positive selection

We applied two statistics to the masked dataset to identify regions of the genome with patterns of genetic variability compatible with positive selection. Within each population, we used Tajima’s D (Tajima, 1989) based on nucleotide diversity and number of segregating sites. Tajima’s D was calculated in 2 Mbp non-overlapping windows using vcftools - TajimaD 2000000. We also used XP-EHH (cross-population extended haplotype homozygosity), a haplotype-based method, which allows the identification of regions under selection by comparing two different populations. XP-EHH was calculated using Selscan v.2 (Szpiech, 2022; Szpiech & Hernandez, 2014) --xpehh with options -- trunc-ok to include calculations at the boundaries of chromosomes, --max-extend 0 for no distance restriction in the calculation of EHH, and default –cutoff 0.5 for EHH decay stopping condition. Then, we searched for 2Mb windows with extreme values using Selscan function --norm with parameters - -crit-percent 0.05 --bp-win --winsize 2000000. Per-chromosome distribution of Tajima’s D and XP-EHH(DSE-DNW) values can be found in Figures S1 and S2.

### Investigating the biological effect of the regions putatively under selection

Windows of interest based on patterns of genetic differentiation were further studied to focus on changes in the biological function and investigate their possible underlying biological cause. Vcftools –bed command was used to extract the windows of interest. The impact on the biological function of polymorphisms falling in the coding regions within these windows was estimated using Ensembl’s Variant Effect Predictor (VEP). Overrepresentation of certain biological pathways was then tested by performing a Gene Ontology enrichment analysis. Genes containing High and Moderate impact variants were tested for statistical overrepresentation in g:Profiler (Raudvere et al., 2019) using all known genes annotated in the Durum Reference genome as background and a significance g:SCS threshold of 0.05 (default).

## Acknowledgments

A.I. has been supported by an FPI fellowship (PRE2018-083529). L. B. is a Ramón y Cajal Fellow. (RYC2018-024770-I) both fellowships funded by the Ministerio de Ciencia e Innovación—Agencia Estatal de Investigación/Fondo Social Europeo. We acknowledge financial support from the Spanish Agencia Estatal de Investigación (Ministry of Science and Innovation-State Research Agency) (AEI), through the “Severo Ochoa Programme for Centres of Excellence in R&D” SEV-2015-0533 and CEX2019-000902-S. This work was also supported by the CERCA programme by the Generalitat de Catalunya. The Authors thank Cristóbal Uauy for his valuable comments on the manuscript.

## Bibliography

Akter, N., & Rafiqul Islam, M. (2017). Heat stress effects and management in wheat. A review. Agronomy for Sustainable Development, 37(5), 37. 10.1007/s13593-017-0443-9

Arora, N. K. (2019). Impact of climate change on agriculture production and its sustainable solutions. Environmental Sustainability, 2(2), 95–96. 10.1007/s42398-019-00078-w

Arranz-Otaegui, A., Colledge, S., Zapata, L., Teira-Mayolini, L. C., & Ibáñez, J. J. (2016). Regional diversity on the timing for the initial appearance of cereal cultivation and domestication in southwest Asia. Proceedings of the National Academy of Sciences of the United States of America, 113(49), 14001–14006. 10.1073/pnas.1612797113

Asouti, E., & Fuller, D. Q. (2013). A Contextual Approach to the Emergence of Agriculture in Southwest Asia: Reconstructing Early Neolithic Plant-Food Production. Current Anthropology, 54(3), 299–345. 10.1086/670679

Avni, R., Nave, M., Barad, O., Baruch, K., Twardziok, S. O., Gundlach, H., Hale, I., Mascher, M., Spannagl, M., Wiebe, K., Jordan, K. W., Golan, G., Deek, J., Ben-Zvi, B., Ben-zvi, G., Himmelbach, A., Maclachlan, R. P., Sharpe, A. G., Komatsuda, T., … Distelfeld, A. (2017). Wild emmer genome architecture and diversity elucidate wheat evolution and domestication. Science, 97(July), 93–97. 10.1126/science.aan0032

Bahaji, A., Li, J., Sánchez-López, Á. M., Baroja-Fernández, E., Muñoz, F. J., Ovecka, M., Almagro, G., Montero, M., Ezquer, I., Etxeberria, E., & Pozueta-Romero, J. (2014). Starch biosynthesis, its regulation and biotechnological approaches to improve crop yields. Biotechnology Advances, 32(1), 87–106. 10.1016/j.biotechadv.2013.06.006

Bukowski, R., Guo, X., Lu, Y., Zou, C., He, B., Rong, Z., Wang, B., Xu, D., Yang, B., Xie, C., Fan, L., Gao, S., Xu, X., Zhang, G., Li, Y., Jiao, Y., Doebley, J. F., Ross-Ibarra, J., Lorant, A., … Xu, Y. (2018). Construction of the third-generation Zea mays haplotype map. GigaScience, 7(4), 1–12. 10.1093/gigascience/gix134

Carmona, D., Lajeunesse, M. J., & Johnson, M. T. J. (2011). Plant traits that predict resistance to herbivores. Functional Ecology, 25(2), 358–367. 10.1111/j.1365-2435.2010.01794.x

Chacón-Duque, J. C., Adhikari, K., Fuentes-Guajardo, M., Mendoza-Revilla, J., Acuña-Alonzo, V., Barquera, R., Quinto-Sánchez, M., Gómez-Valdés, J., Everardo Martínez, P., Villamil-Ramírez, H., Hünemeier, T., Ramallo, V., Silva de Cerqueira, C. C., Hurtado, M., Villegas, V., Granja, V., Villena, M., Vásquez, R., Llop, E., … Ruiz-Linares, A. (2018). Latin Americans show wide-spread Converso ancestry and imprint of local Native ancestry on physical appearance. Nature Communications, 9(1). 10.1038/s41467-018-07748-z

Chaudhary, B. (2013). Plant domestication and resistance to herbivory. International Journal of Plant Genomics, 2013. 10.1155/2013/572784

Chen, H., Zhang, D., Guo, J., Wu, H., Jin, M., Lu, Q., Lu, C., & Zhang, L. (2006). A Psb27 homologue in Arabidopsis thaliana is required for efficient repair of photodamaged photosystem II. Plant Molecular Biology, 61(4), 567–575. 10.1007/s11103-006-0031-x

Chen, Y. H., Gols, R., & Benrey, B. (2015). Crop Domestication and Its Impact on Naturally Selected Trophic Interactions. Annual Review of Entomology, 60(1), 35–58. 10.1146/annurev-ento-010814-020601

Cheng, H., Liu, J., Wen, J., Nie, X., Xu, L., Chen, N., Li, Z., Wang, Q., Zheng, Z., Li, M., Cui, L., Liu, Z., Bian, J., Wang, Z., Xu, S., Yang, Q., Appels, R., Han, D., Song, W., … Jiang, Y. (2019). Frequent intra- and inter-species introgression shapes the landscape of genetic variation in bread wheat. Genome Biology, 20(1), 1–16. 10.1186/s13059-019-1744-x

Civáň, P., Ivaničová, Z., & Brown, T. A. (2013). Reticulated origin of domesticated emmer wheat supports a dynamic model for the emergence of agriculture in the fertile crescent. PLoS ONE, 8(11), 1–11. 10.1371/journal.pone.0081955

Clark, M. A., Domingo, N. G. G., Colgan, K., Thakrar, S. K., Tilman, D., Lynch, J., Azevedo, I. L., & Hill, J. D. (2020). Global food system emissions could preclude achieving the 1.5° and 2°C climate change targets. Science, 370(6517), 705–708. 10.1126/science.aba7357

Cook, D. E. (n.d.). generate masked ranges.

Cortés, A. J., & López-Hernández, F. (2021). Harnessing crop wild diversity for climate change adaptation. Genes, 12(5), NA. 10.3390/genes12050783

Danecek, P., Bonfield, J. K., Liddle, J., Marshall, J., Ohan, V., Pollard, M. O., Whitwham, A., Keane, T., McCarthy, S. A., Davies, R. M., & Li, H. (2021). Twelve years of SAMtools and BCFtools. GigaScience, 10(2), 1–4. 10.1093/gigascience/giab008

Delaneau, O., Marchini, J., McVeanh, G. A., Donnelly, P., Lunter, G., Marchini, J. L., Myers, S., Gupta-Hinch, A., Iqbal, Z., Mathieson, I., Rimmer, A., Xifara, D. K., Kerasidou, A., Churchhouse, C., Altshuler, D. M., Gabriel, S. B., Lander, E. S., Gupta, N., Daly, M. J., … Peltonen, L. (2014). Integrating sequence and array data to create an improved 1000 Genomes Project haplotype reference panel. Nature Communications, 5, 1–9. 10.1038/ncomms4934

Durrant, W. E., & Dong, X. (2004). SYSTEMIC ACQUIRED RESISTANCE. Annual Review of Phytopathology, 42(1), 185–209. 10.1146/annurev.phyto.42.040803.140421

Dvořák, J. (2001). Triticum Species (Wheat). Encyclopedia of Genetics, 2060–2068. 10.1006/rwgn.2001.1672

Engels, J. M. M., & Thormann, I. (2020). Main Challenges and Actions Needed to Improve Conservation and Sustainable Use of Our Crop Wild Relatives. *Plants (Basel*, Switzerland), 9(8). 10.3390/plants9080968

Eydivandi, S., Roudbar, M. A., Karimi, M. O., & Sahana, G. (2021). Genomic scans for selective sweeps through haplotype homozygosity and allelic fixation in 14 indigenous sheep breeds from Middle East and South Asia. Scientific Reports, 11(1), 2834. 10.1038/s41598-021-82625-2

FAO. (2021). Agricultural production statistics 2000-2021. In FAOSTAT ANALYTICAL BRIEF 60. https://www.fao.org/3/cc3751en/cc3751en.pdf

Finkina, E. I., Melnikova, D. N., Bogdanov, I. V, & Ovchinnikova, T. V. (2016). Lipid Transfer Proteins As Components of the Plant Innate Immune System: Structure, Functions, and Applications. Acta Naturae, 8(2), 47–61.

Fraire-Velazquez, S., & Emmanuel, V. (2013). Abiotic Stress in Plants and Metabolic Responses. Abiotic Stress - Plant Responses and Applications in Agriculture. 10.5772/54859

Fuller, D. Q., Denham, T., & Allaby, R. (2023). Plant domestication and agricultural ecologies. Current Biology, 33. 10.1016/j.cub.2023.04.038

Golenberg, E. M. (1988). Outcrossing rates and their relationship to phenology in Triticum dicoccoides . Theoretical and Applied Genetics, 75, 937–944. 10.1007/BF00258057

Gonthier, D. J., Ennis, K. K., Farinas, S., Hsieh, H.-Y., Iverson, A. L., Batáry, P., Rudolphi, J., Tscharntke, T., Cardinale, B. J., & Perfecto, I. (2014). Biodiversity conservation in agriculture requires a multi-scale approach. *Proceedings*. Biological Sciences, 281(1791), 20141358. 10.1098/rspb.2014.1358

Gutaker, R. M., Zaidem, M., Fu, Y.-B., Diederichsen, A., Smith, O., Ware, R., & Allaby, R. G. (2019). Flax latitudinal adaptation at LuTFL1 altered architecture and promoted fiber production. Scientific Reports, 9(1), 976. 10.1038/s41598-018-37086-5

Hajjar, R., & Hodgkin, T. (2007). The use of wild relatives in crop improvement: a survey of developments over the last 20 years. Euphytica, 156(1), 1–13. 10.1007/s10681-007-9363-0

Hazen, S. P., Hawley, R. M., Davis, G. L., Henrissat, B., & Walton, J. D. (2003). Quantitative Trait Loci and Comparative Genomics of Cereal Cell Wall Composition. Plant Physiology, 132(1), 263–271. 10.1104/pp.103.020016

Hellenthal, G., Busby, G. B. J., Band, G., Wilson, J. F., Capelli, C., Falush, D., & Myers, S. (2014). A genetic atlas of human admixture history. Science, 343(6172), 747–751. 10.1126/science.1243518

Henderson, E. C., & Brelsford, A. (2020). Genomic differentiation across the speciation continuum in three hummingbird species pairs. BMC Evolutionary Biology, 20(1), 1–11. 10.1186/s12862-020-01674-9

Hu, S., Ding, Y., & Zhu, C. (2020). Sensitivity and Responses of Chloroplasts to Heat Stress in Plants. Frontiers in Plant Science, 11(April), 1–11. 10.3389/fpls.2020.00375

Hufford, M. B., Berny Mier Y Teran, J. C., & Gepts, P. (2019). Crop Biodiversity: An Unfinished Magnum Opus of Nature. Annual Review of Plant Biology, 70, 727–751. 10.1146/annurev-arplant-042817-040240

Iob, A., & Botigué, L. (2022). Genomic analysis of emmer wheat shows a complex history with two distinct domestic groups and evidence of differential hybridization with wild emmer from the western Fertile Crescent. Vegetation History and Archaeobotany, 0123456789. 10.1007/s00334-022-00898-7

Janzen, G. M., Wang, L., & Hufford, M. B. (2019). The extent of adaptive wild introgression in crops. New Phytologist, 221(3), 1279–1288. 10.1111/nph.15457

Jiang, Y. F., Lan, X. J., Luo, W., Kong, X. C., Qi, P. F., Wang, J. R., Wei, Y. M., Jiang, Q. T., Liu, Y. X., Peng, Y. Y., Chen, G. Y., Dai, S. F., & Zheng, Y. L. (2014). Genome-wide quantitative trait locus mapping identifies multiple major loci for brittle rachis and threshability in Tibetan semi-wild wheat (Triticum aestivum ssp. tibetanum Shao). PLoS ONE, 9(12), 1–15. 10.1371/journal.pone.0114066

Johnson, V. M., Biswas, S., Roose, J. L., Pakrasi, H. B., & Liu, H. (2022). Psb27, a photosystem II assembly protein, enables quenching of excess light energy during its participation in the PSII lifecycle. Photosynthesis Research, 152(3), 297–304. 10.1007/s11120-021-00895-3

Khoury, C. K., Brush, S., Costich, D. E., Curry, H. A., de Haan, S., Engels, J. M. M., Guarino, L., Hoban, S., Mercer, K. L., Miller, A. J., Nabhan, G. P., Perales, H. R., Richards, C., Riggins, C., & Thormann, I. (2022). Crop genetic erosion: understanding and responding to loss of crop diversity. New Phytologist, 233(1), 84–118. 10.1111/nph.17733

Labeyrie, V., Renard, D., Aumeeruddy-Thomas, Y., Benyei, P., Caillon, S., Calvet-Mir, L., M. Carrière, S., Demongeot, M., Descamps, E., Braga Junqueira, A., Li, X., Locqueville, J., Mattalia, G., Miñarro, S., Morel, A., Porcuna-Ferrer, A., Schlingmann, A., Vieira da Cunha Avila, J., & Reyes-García, V. (2021). The role of crop diversity in climate change adaptation: insights from local observations to inform decision making in agriculture. Current Opinion in Environmental Sustainability, 51, 15–23. 10.1016/j.cosust.2021.01.006

Lawson, D. J., Hellenthal, G., Myers, S., & Falush, D. (2012). Inference of population structure using dense haplotype data. PLoS Genetics, 8(1), 11–17. 10.1371/journal.pgen.1002453

Li, H., & Wren, J. (2014). Toward better understanding of artifacts in variant calling from high-coverage samples. Bioinformatics, 30(20), 2843–2851. 10.1093/bioinformatics/btu356

Liu, H., Huang, R. Y. C., Chen, J., Gross, M. L., & Pakrasi, H. B. (2011). Psb27, a transiently associated protein, binds to the chlorophyll binding protein CP43 in photosystem II assembly intermediates. Proceedings of the National Academy of Sciences of the United States of America, 108(45), 18536–18541. 10.1073/pnas.1111597108

Lucas, S. J., Salantur, A., Yazar, S., & Budak, H. (2017). High-throughput SNP genotyping of modern and wild emmer wheat for yield and root morphology using a combined association and linkage analysis. Functional & Integrative Genomics, 17(6), 667–685. 10.1007/s10142-017-0563-y

Luo, M. C., Yang, Z. L., You, F. M., Kawahara, T., Waines, J. G., & Dvorak, J. (2007). The structure of wild and domesticated emmer wheat populations, gene flow between them, and the site of emmer domestication. Theoretical and Applied Genetics, 114(6), 947–959. 10.1007/s00122-006-0474-0

Lv, L., Zhang, W., Sun, L., Zhao, A., Zhang, Y., Wang, L., Liu, Y., Li, Z., Li, H., & Chen, X. (2020). Gene co-expression network analysis to identify critical modules and candidate genes of drought-resistance in wheat. PLOS ONE, 15(8), e0236186. 10.1371/journal.pone.0236186

Maccaferri, M., Harris, N. S., Twardziok, S. O., Pasam, R. K., Gundlach, H., Spannagl, M., Ormanbekova, D., Lux, T., Prade, V. M., Milner, S. G., Himmelbach, A., Mascher, M., Bagnaresi, P., Faccioli, P., Cozzi, P., Lauria, M., Lazzari, B., Stella, A., Manconi, A., … Cattivelli, L. (2019). Durum wheat genome highlights past domestication signatures and future improvement targets. Nature Genetics, 51(5), 885–895. 10.1038/s41588-019-0381-3

Marone, D., Russo, M. A., Mores, A., Ficco, D. B. M., Laidò, G., Mastrangelo, A. M., & Borrelli, G. M. (2021). Importance of landraces in cereal breeding for stress tolerance. Plants, 10(7). 10.3390/plants10071267

Martin, S. (n.d.). genomics general.

Mithen, S., Richardson, A., & Finlayson, B. (2023). The flow of ideas: shared symbolism during the Neolithic emergence in Southwest Asia: WF16 and Göbekli Tepe. Antiquity, October 2022, 1–21. 10.15184/aqy.2023.67

Mohammadi, M., Mirlohi, A., Majidi, M. M., & Soleimani Kartalaei, E. (2021). Emmer wheat as a source for trait improvement in durum wheat: a study of general and specific combining ability. Euphytica, 217(4), 1–20. 10.1007/s10681-021-02796-x

Nave, M., Avni, R., Çakır, E., Portnoy, V., Sela, H., Pourkheirandish, M., Ozkan, H., Hale, I., Komatsuda, T., Dvorak, J., & Distelfeld, A. (2019). Wheat domestication in light of haplotype analyses of the Brittle rachis 1 genes (BTR1-A and BTR1-B). Plant Science, 285(May), 193–199. 10.1016/j.plantsci.2019.05.012

Oliveira, H. R., Jacocks, L., Czajkowska, B. I., Kennedy, S. L., & Brown, T. A. (2020). Multiregional origins of the domesticated tetraploid wheats. PLoS ONE, 15(1), 1–20. 10.1371/journal.pone.0227148

Ortiz-Bobea, A., Ault, T. R. T. R., Carrillo, C. M. C. M., Chambers, R. G., & Lobell, D. B. (2021). Anthropogenic climate change has slowed global agricultural productivity growth. Nature Climate Change, 11(4), 306–312. 10.1038/s41558-021-01000-1

Ozkan, H., Brandolini, A., Schafer-Pregl, R., & Salamini, F. (2002). AFLP analysis of a collection of tetraploid wheat indicated the origin of emmer and hard wheat domestication in south-eastern Turkey. Molecular Biology and Evolution, 19(10), 1797–1801. 10.1093/oxfordjournals.molbev.a004002

Ozkan, H., Willcox, G., Graner, A., Salamini, F., & Kilian, B. (2011). Geographic distribution and domestication of wild emmer wheat (Triticum dicoccoides). Genetic Resources and Crop Evolution, 58, 11–53. 10.1007/s10722-010-9581-5

Özkan, H., Willcox, G., Graner, A., Salamini, F., Kilian, B., Hakan Ozkan • George Willcox • Andreas Graner • Francesco Salamini • Benjamin Kilian, & Received: (2011). Geographic distribution and domestication of wild emmer wheat (Triticum dicoccoides). Genetic Resources and Crop Evolution, 58(1), 11–53. 10.1007/s10722-010-9581-5

Pont, C., Leroy, T., Seidel, M., & Tondelli, A. (2019). Tracing the ancestry of modern bread wheats. Nature Genetics, 51, 905–911. 10.1038/s41588-019-0393-z Tracing

Purcell, S., Neale, B., Todd-Brown, K., Thomas, L., Ferreira, M., Bender, D., Maller, J., Sklar, P., de Bakker, P., Daly, M., & Sham, P. (2007). PLINK: a toolset for whole-genome association and population-based linkage analysis. American Journal of Human Genetics, 81.

Qian, Y., Meng, M., Zhou, C., Liu, H., Jiang, H., Xu, Y., Chen, W., Ding, Z., Liu, Y., Gong, X., Wang, C., Lei, Y., Wang, T., Wang, Y., Gan, X., Meyer, A., He, S., & Yang, L. (2023). The Role of Introgression During the Radiation of Endemic Fishes Adapted to Living at Extreme Altitudes in the Tibetan Plateau. Molecular Biology and Evolution, 40(6), msad129. 10.1093/molbev/msad129

R Core Team. (2021). R: A language and environment for statistical computing. R Foundation for Statistical Computing, Vienna, Austria. https://www.r-project.org/

Raudvere, U., Kolberg, L., Kuzmin, I., Arak, T., Adler, P., Peterson, H., & Vilo, J. (2019). G:Profiler: A web server for functional enrichment analysis and conversions of gene lists (2019 update). Nucleic Acids Research, 47(W1), W191–W198. 10.1093/nar/gkz369

Reiter, W.-D. (2002). Biosynthesis and properties of the plant cell wall. Current Opinion in Plant Biology, 5(6), 536–542. 10.1016/S1369-5266(02)00306-0

Saleh, M. M. (2020). Stress breeding of neglected tetraploid primitive wheat (Triticum dicoccum, Triticum carthlicum and Triticum polonicum). Current Botany, 11, 99–110. 10.25081/cb.2020.v11.6100

Scott, M. F., Botigué, L. R., Brace, S., Stevens, C. J., Mullin, V. E., Stevenson, A., Thomas, M. G., Fuller, D. Q., & Mott, R. (2019). A 3,000-year-old Egyptian emmer wheat genome reveals dispersal and domestication history. Nature Plants, 5(11), 1120–1128. 10.1038/s41477-019-0534-5

Simons, K. J., Fellers, J. P., Trick, H. N., Zhang, Z., Tai, Y. S., Gill, B. S., & Faris, J. D. (2006). Molecular characterization of the major wheat domestication gene Q. Genetics, 172(1), 547–555. 10.1534/genetics.105.044727

Singh, J., & van der Knaap, E. (2022). Unintended Consequences of Plant Domestication. Plant and Cell Physiology, 63(11), 1573–1583. 10.1093/pcp/pcac083

Streit Krug, A., Drummond, E. B. M., Van Tassel, D. L., & Warschefsky, E. J. (2023). The next era of crop domestication starts now. Proceedings of the National Academy of Sciences, 120. 10.1073/pnas.2205769120

Swarup, S., Cargill, E. J., Crosby, K., Flagel, L., Kniskern, J., & Glenn, K. C. (2021). Genetic diversity is indispensable for plant breeding to improve crops. Crop Science, 61(2), 839–852. 10.1002/csc2.20377

Szpiech, Z. A. (2022). Selscan 2.0: Scanning for Sweeps in Unphased Data. BioRxiv, 2021.10.22.465497.

Szpiech, Z. A., & Hernandez, R. D. (2014). Selscan: An efficient multithreaded program to perform EHH-based scans for positive selection. Molecular Biology and Evolution, 31(10), 2824–2827. 10.1093/molbev/msu211

Tajima, F. (1989). Statistical method for testing the neutral mutation hypothesis by DNA polymorphism. Genetics, 123(3), 585–595. 10.1093/genetics/123.3.585

Tetlow, I. J., & Emes, M. J. (2014). A review of starch-branching enzymes and their role in amylopectin biosynthesis. IUBMB Life, 66(8), 546–558. 10.1002/iub.1297

Tryfona, T., Theys, T. E., Wagner, T., Stott, K., Keegstra, K., & Dupree, P. (2014). Characterisation of FUT4 and FUT6 α-(1 → 2)-fucosyltransferases reveals that absence of root arabinogalactan fucosylation increases Arabidopsis root growth salt sensitivity. PloS One, 9(3), e93291. 10.1371/journal.pone.0093291

Van der Auwera, GA O’Connor, B. (2020). Genomics in the Cloud: Using Docker, GATK, and WDL in Terra (1st Edition).

Venkateswaran, K., Elangovan, M., & Sivaraj, N. (2019). Chapter 2 - Origin, Domestication and Diffusion of Sorghum bicolor. In C. Aruna, K. B. R. S. Visarada, B. V. Bhat, & V. A. B. T.-B. S. for D. E. U. Tonapi (Eds.), Woodhead Publishing Series in Food Science, Technology and Nutrition (pp. 15–31). Woodhead Publishing. 10.1016/B978-0-08-101879-8.00002-4

Wang, Q. L., Chen, J. H., He, N. Y., & Guo, F. Q. (2018). Metabolic reprogramming in chloroplasts under heat stress in plants. International Journal of Molecular Sciences, 19(3), 9–11. 10.3390/ijms19030849

Wang, Z., Wang, W., Xie, X., Wang, Y., Yang, Z., Peng, H., Xin, M., Yao, Y., Hu, Z., Liu, J., Su, Z., Xie, C., Li, B., Ni, Z., Sun, Q., & Guo, W. (2022). Dispersed emergence and protracted domestication of polyploid wheat uncovered by mosaic ancestral haploblock inference. Nature Communications, 13(2022), 1–14. 10.1038/s41467-022-31581-0

Wangkumhang, P., Greenfield, M., & Hellenthal, G. (2022). An efficient method to identify, date, and describe admixture events using haplotype information. Genome Research, 32(8), 1553– 1564. 10.1101/gr.275994.121

Xu, J., Yuan, Y., Xu, Y., Zhang, G., Guo, X., Wu, F., Wang, Q., Rong, T., Pan, G., Cao, M., Tang, Q., Gao, S., Liu, Y., Wang, J., Lan, H., & Lu, Y. (2014). Identification of candidate genes for drought tolerance by whole-genome resequencing in maize. BMC Plant Biology, 14(1), 83. 10.1186/1471-2229-14-83

Zaharieva, M., Ayana, N. G., Hakimi, A. Al, Misra, S. C., Monneveux, P., Ayana, N. G., Hakimi, A. Al, Zaharieva, M., Ayana, N. G., Hakimi, A. Al, Misra, S. C., Monneveux, P., Ayana, N. G., Hakimi, A. Al, Zaharieva, M., Ayana, N. G., Hakimi, A. Al, Misra, S. C., & Monneveux, P. (2010). Cultivated emmer wheat (Triticum dicoccon Schrank), an old crop with promising future: A review. Genetic Resources and Crop Evolution, 57(6), 937–962. 10.1007/s10722-010-9572-6

Zhang, L., Paasch, B. C., Chen, J., Day, B., & He, S. Y. (2019). An important role of l-fucose biosynthesis and protein fucosylation genes in Arabidopsis immunity. The New Phytologist, 222(2), 981–994. 10.1111/nph.15639

Zhao, C., Liu, B., Piao, S., Wang, X., Lobell, D. B., Huang, Y., Huang, M., Yao, Y., Bassu, S., Ciais, P., Durand, J. L., Elliott, J., Ewert, F., Janssens, I. A., Li, T., Lin, E., Liu, Q., Martre, P., Müller, C., … Asseng, S. (2017). Temperature increase reduces global yields of major crops in four independent estimates. Proceedings of the National Academy of Sciences of the United States of America, 114(35), 9326–9331. 10.1073/pnas.1701762114

Zhao, X., Guo, Y., Kang, L., Yin, C., Bi, A., Xu, D., Zhang, Z., Zhang, J., Yang, X., Xu, J., Xu, S., Song, X., Zhang, M., Li, Y., Kear, P., Wang, J., Liu, Z., Fu, X., & Lu, F. (2023). Population genomics unravels the Holocene history of bread wheat and its relatives. Nature Plants, 9(March). 10.1038/s41477-023-01367-3

Zhou, Y., Zhao, X., Li, Y., Xu, J., Bi, A., Kang, L., Xu, D., Chen, H., Wang, Y., Wang, Y. ge, Liu, S., Jiao, C., Lu, H., Wang, J., Yin, C., Jiao, Y., & Lu, F. (2020). Triticum population sequencing provides insights into wheat adaptation. Nature Genetics, 52(12), 1412–1422. 10.1038/s41588-020-00722-w

Zohary, D. (2013). Domestication of Crop Plants. *Encyclopedia of Biodiversity: Second Edition*, March 2018, 657–664. 10.1016/B978-0-12-384719-5.00199-4

